# Improved CD4 T-cell profile and inflammatory levels in HIV-infected subjects on maraviroc-containing therapy is associated with better responsiveness to HBV vaccination

**DOI:** 10.1101/298521

**Authors:** Inés Herrero-Fernéndez, Isaac Rosado-Sánchez, Miguel Genebat, Laura Tarancón-Díez, María Mar Rodríguez-Méndez, María Mar Pozo-Balado, Carmen Lozano, Ezequiel Ruiz-Mateos, Manuel Leal, Yolanda M. Pacheco

## Abstract

**Introduction:** We previously found that a maraviroc-containing combined antiretroviral therapy (MVC-cART) was associated with a better response to the Hepatitis B Virus (HBV) vaccine in HIV-infected subjects younger than 50 years old. We aimed here to extend our previous analysis including immunological parameters related to inflammation, T-cell and dendritic cell (DC) subsets phenotype and to explore the impact of MVC-cART on these parameters.

**Methods:** We analyzed baseline samples of vaccinated subjects under 50 years old (n=41). We characterized CD4 T-cells according to the distribution of their maturational subsets and the expression of activation, senescence and prone-to-apoptosis markers; we also quantified Treg-cells and main DC subsets. Linear regressions were performed to determine the impact of these variables on the magnitude of vaccine response. Binary logistic regressions were explored to analyze the impact of MVC-cART on immunological parameters. Correlations with the time of MVC exposure were also explored.

**Results:** MVC-cART remained independently associated with HBV-vaccine responsiveness even after adjusting by immunological variables. The %CD4^+^CD25^hi^FoxP3^+^ki67^+^ and %pDCs were also independently associated. Moreover, HIV-infected subjects on MVC-containing therapy prior to vaccination showed lower inflammatory levels, activated CD4 T-cells and frequency of Treg cells and higher frequency of recent thymic emigrants.

**Conclusion:** Treg-cell levels negatively impacted the HBV-vaccine response, whereas higher pDCs levels and a MVC-cART prior to vaccination were associated with better responsiveness. These factors together with the improved phenotypic CD4 T-cell profile and the lower inflammatory levels found in subjects with a MVC-cART prior HBV vaccination could contribute to their enhanced vaccine response.

## INTRODUCTION

Human Inmunodeficiency Virus (HIV)-infected subjects are at high risk for Hepatitis B Virus (HBV) infection and progression of severe, life-threatening hepatic complications, as cirrhosis and hepatocellular carcinoma [1, 2]. To prevent the associated morbimortality, worldwide current guidelines recommend vaccination against HBV in all HIV-infected subjects susceptible to be coinfected by HBV, but the response rates are lower than in HIV non-infected subjects [reviewed in 3].

The best known predictors of vaccine efficacy are undetectable viral load and CD4 T-cell counts above 350 cells/mm^3^ [3]. Thus, it is well assumed that a successful combined antiretroviral therapy (cART) favors the vaccine response, however, the influence of the type of antiretroviral treatment has been scarcely explored until now. In this line, it was first described that maraviroc (MVC), a CCR5 antagonist, enhanced meningococcal neo-immunization and accelerated the response to tetanus boost [4]. More recently, we have also reported that a MVC-containing cART (MVC-cART) was associated with a better response against the HBV vaccine, at least in subjects younger than 50 years old [5]. Nevertheless, the potential subjacent mechanisms were unaddressed.

Different antiretroviral combinations including MVC, have comparatively proved their beneficial effects on the levels of inflammatory biomarkers [6] and the T-cell immunophenotype [7]. In two clinical trials, an improvement of duodenal immunity and a reduction in bone loss has been associated to such combinations [8, 9]. Furthermore, MVC on monotherapy also reduced the frequency of regulatory T-cells (Treg) in antiretroviral-naïve subjects [10], even improving the distribution of Treg subsets [11]. This could be relevant since we observed that regulatory T cells (Treg) negatively impacted the HBV vaccine responsiveness in a previous cohort [12]. Possibly, MVC could enhance different functions required to mount an effective response following HBV vaccination including antigen-presentation, T-cell help, regulatory T-cell suppression and B cell functions [13, 14].

In the present work, we extended our previous analysis [5] including several relevant immunological variables to gain knowledge about the factors impacting the responsiveness to the HBV vaccine. We additionally explored the potential effect of a MVC-cART exposure on different parameters related to inflammation, T-cell function and DC subsets that could account for the effectiveness of the HBV vaccination.

## RESULTS

### Demographic, clinical and immunological variables associated with the magnitude of HBV vaccine responsiveness

We extended our previous analysis including demographic and clinical variables [5], adding several relevant immunological variables to explore to what extent each of these factors could impact the magnitude of the response. Around half of the population (51%) was receiving a MVC-cART, consisting of MVC and a boosted protease inhibitor (PI) or MVC and two nucleoside-reverse transcriptase inhibitors (NRTIs).

As shown in Table 1, sex, previous AIDS, MVC-containing-cART, previous HBV vaccination, simultaneous HAV vaccination, hsCRP, %CD4^+^ki67^+^, %CD4^+^CD25^hi^FoxP3^+^HLA-DR^+^, %CD4^+^CD25^hi^FoxP3^+^ki67^+^ and %pDCs showed a *p* value <0.1 in the univariate analyses, thus, were all introduced in the multivariate model. In this model, the unique variable showing independent association was MVC-containing-cART, with a borderline significance (*p*=0.059; B [95% CI], 243.4 [-10.2-504.9]). Next, we explored a stepwise forward multivariate analysis. In this model, the only variables included were MVC-containing-cART (*p*=0.016; B [95% CI], 293.3 [58.1-5280.5]), %CD4^+^CD25^hi^FoxP3^+^ki67^+^ (*p*=0.005, B [95% CI], -29.5 [-49.6—9.5]) and %pDCs (p=0.009, B [95% CI], 1961.2 [530.4-3392.1]) (Table 1), which were independently associated with the magnitude of response. Interestingly, the association was negative with the %CD4^+^CD25^hi^FoxP3^+^ki67^+^ but positive with the MVC-containing-cART and %pDCs.

**Table 1.** Demographic, clinical and immunological variables associated with HBV vaccine responsiveness

| Clinical and immunonological variables | n=41 | Unadjusted P value;<br>B (95% CI) | Adjusted P value;<br>B (95% CI) |
| --- | --- | --- | --- |
| Male sex, n (%) | 31 (76) | **0.055; -294.2 [-595.5-7.1]** |  |
| Age (years) | 41.0 [33.5-46.5] | 0.717; -3.3 [-21.3-14.8] |  |
| Nadir CD4<sup>+</sup> T-cell count (cells/mm<sup>3</sup>) | 287.0 [206.3-402.0] | 0.288; 0.46 [-0.40-1.32] |  |
| CD4<sup>+</sup> T-cell count (cells/mm<sup>3</sup>) | 703.0 [553.0-887.5] | 0.302; 0.30 [-0.28-0.88] |  |
| CD8<sup>+</sup> T-cell count (cells/mm<sup>3</sup>) | 643.0 [501.0-927.0] | 0.344; -0.19 [-0.58-0.21] |  |
| Ratio CD4<sup>+</sup>/CD8<sup>+</sup> | 0.9 [0.8-1.4] | 0.205; 139.1 [-79.0-357.1] |  |
| Time since diagnosis (months) | 75.0 [37.5-212.0] | 0.200; 0.89 [-0.49-2.27] |  |
| Sexual transmission, n (%) | 36 (88) | 0.579; 114.4 [-298.7-527.4] |  |
| Previous AIDS, n (%) | 2 (5) | **0.074; -548.2 [-1152.6-56.3]** |  |
| Previous HCV coinfection, n (%) | 5 (12) | 0.453; 154.4 [-257.3-566.0] |  |
| NRTI containing cART, n (%) | 18 (44) | 0.496; 92.4 [-179.4-364.2] |  |
| MVC containing cART, n (%) | 21 (51) | **0.017; 310.8 [58.7-563.0]** | **0.016; 293.3 [58.1-5280.5]** |
| Previous HBV vaccination, n (%)* | 13 (32) | **0.067; 262.6 [-19.5-544.8]** |  |
| Simultaneous HAV vaccination, n (%) | 18 (44) | **0.029; 287.9 [31.0-545.0]** |  |
| Time on HIV-treatment (months) | 69.5 [38.0-178.8] | 0.376; 0.74 [-0.93-2.42] |  |
| hsCRP (mg/L) | 0.8 [0.5-1.3] | **0.096; -165.8 [-362.4-30.9]** |  |
| % Lymphocytes | 35.6 [26.1-39.2] | 0.555; 4.9 [-11.7-21.4] |  |
| % Monocytes | 5.6 [5.3-6.1] | 0.393; 66.5 [-89.5-222.4] |  |
| % Neutrophils | 53.6 [49.3-63.3] | 0.593; -3.7 [-17.8-10.3] |  |
| % CD4<sup>+</sup> Naives | 44.6 [34.2-51.0] | 0.496; 3.2 [-6.3-12.7] |  |
| % CD4<sup>+</sup> RTE | 75.6 [66.5-79.8] | 0.204; 9.2 [-5.2-23.7] |  |
| % CD4<sup>+</sup> Central Memory | 28.3 [23.2-39.5] | 0.791; -1.9 [-16. 7-12. 8] |  |
| % CD4<sup>+</sup> Effector Memory | 20.7 [14.9-27.8] | 0.400; -6.1 [-20.5-8.4] |  |
| % CD4<sup>+</sup> TemRA | 1.8 [1.1-4.3] | 0.154; -31.4 [-75.2-12.3] |  |
| % CD4<sup>+</sup> HLA-DR<sup>+</sup> | 1.7 [1.1-2.0] | 0.161; -128.1 [-309.6-53.4] |  |
| % CD4<sup>+</sup> ki67<sup>+</sup> | 2.3 [2.1-2.7] | **0.053; -199.5 [-401.4-2.5]** |  |
| % CD4<sup>+</sup> CD57<sup>+</sup> | 4.7 [3.1-8.0] | 0.825; -3.1 [-31.3-25.2] |  |
| % CD4<sup>+</sup> CD95<sup>+</sup> | 55.0 [43.1-65.0] | 0.394; -3.7 [-12.3-5.0] |  |
| % CD4<sup>+</sup> CD25<sup>hi</sup>FoxP3<sup>+</sup> | 1.3 [1.0-1.8] | 0.565; -64.7 [-290.3-160.9] |  |
| % CD4<sup>+</sup> CD25<sup>hi</sup>FoxP3<sup>+</sup>HLA-DR<sup>+</sup> | 15.1 [11.0-21.6] | **0.039; -19.7 [-38.3--1.0]** |  |
| % CD4<sup>+</sup> CD25<sup>hi</sup>FoxP3<sup>+</sup>ki67<sup>+</sup> | 19.8 [16.9-25.0] | **0.011; -28.9 [-50.7--7.1]** | **0.005; -29.6 [-49.6--9.5]** |
| % CD4<sup>+</sup> CD25<sup>hi</sup>FoxP3<sup>+</sup>CD39<sup>+</sup> | 81.2 [41.7-87.0] | 0.274; -2.9 [-8.2-2.4] |  |
| % CD4<sup>+</sup> CD25<sup>hi</sup>FoxP3<sup>+</sup>CTLA4<sup>+</sup> | 58.7 [46.9-68.3] | 0.232; -5.9 [-15.8-4.0] |  |
| % mDCs | 0.7 [0.4-1.1] | 0.477; -85.4 [-325.9-155.1] |  |
| % pDCs | 0.2 [0.18-0.24] | **0.072; 1323.5 [-122.5-2769.6]** | **0.009; 1961.2 [530.4-3392.0]** |

Continuous variables are expressed as median values [IQR] and categorical variables are expressed as number of cases (&). All demographic, clinical andimmunological variables with a *p* value of <0.1 in the unadjusted model were included in the adjusted model and are shown in bold. Hence, male sex, previous AIDS, MVC containing cART, previous HBV vaccination, simultaneous HAV vaccination, hsCRP, &CD4^+^ki67^+^, &CD4^+^CD25^hi^FoxP3^+^HLA-DR^+^, &CD4^+^CD25^hi^FoxP3^+^ki67^+^, &pDCs were included in the stepwise forward multivariate model. As shown, the resulting model only retained the variables MVC containing cART, &CD4^+^CD25^hi^FoxP3^+^ki67^+^ and &pDCs. Variables with a *p* value of <0.05 in the adjusted model were considered statistically significant and are shown in bold.

### Demographic, clinical and immunological variables associated with a MVC-containing cART

We compared the demographic, clinical and immunological variables, at the moment of vaccination, between patients receiving a MVC-cART or a MVC-sparing-cART (Table 2). The age, CD4^+^/CD8^+^ ratio, time on HIV-treatment, %CD4^+^RTE and %mDCs showed a *p* value <0.1 in the univariate analyses and were, therefore, included in the multivariate analysis. To note, 19% of the subjects treated with MVC-cART were also receiving NRTIs, whereas the 70% of the subjects on a MVC-sparing-cART were receiving NRTIs (Table 2). Thus, the absence of NRTIs was highly colineal with the presence of MVC and was not included for adjustment. As it is shown, the %CD4^+^RTE (*p*=0.024; OR [95% CI], 1.20 [1.02-1.41]) and %mDCs (*p*=0.048; OR [95% CI], 0.16 [0.02-0.98]) were independently associated with a MVC-cART, whereas CD4^+^/CD8^+^ ratio (*p*=0.086; OR [95% CI], 0.19 [0.03-1.26]), showed association with borderline significance.

**Table 2.** Demographic, clinical and immunological variables associated with an MVC-containing cART.

| Clinical and immunonological variables (n=41) | MVC-containing cART (N=21) | MVC-sparing cART (N=20) | Unadjusted *p* value; OR [95% CI] | Adjusted *p* value; OR [95% CI] |
| --- | --- | --- | --- | --- |
| Male sex, n (%) | 15 (71) | 16 (80) | 0.525; 0.625 [0.147-2.659] |  |
| Age (years) | 36 [31-44] | 44 [39-48] | **0.024; 0.892 [0.800-0.985]** | 0.878; 1.01 [0.87-1.18] |
| Nadir CD4<sup>+</sup> T-cell count (cells/mm<sup>3</sup>) | 294 [184-412] | 264 [206-379] | 0.539; 1.001 [0.997-1.005] |  |
| CD4<sup>+</sup> T-cell count (cells/mm<sup>3</sup>) | 703 [565-869] | 725 [529-911] | 0.775; 1.000 [0.997-1.002] |  |
| CD8<sup>+</sup> T-cell count (cells/mm<sup>3</sup>) | 781 [538-961] | 596 [491-829] | 0.109; 1.002 [1.000-1.004] |  |
| CD4<sup>+</sup>/CD8<sup>+</sup> Ratio | 0.9 [0.6-1.2] | 1.1 [0.9-1.6] | **0.084; 0.309 [0.082-1.170]** | *0.086; 0.20 [0.03-1.26]* |
| Time since diagnosis (months) | 65 [32-212] | 132 [67-235] | 0.158; 0.995 [0.989-1.002] |  |
| Time on HIV-treatment (months) | 46 [33-147] | 120 [64-201] | **0.057; 0.992 [0.984-1.000]** | 0.105; 0.99 [0.97-1.00] |
| Sexual transmission. n (%) | 18 (86) | 18 (90) | 0.677; 1.500 [0.223-10.077] |  |
| Previous AIDS. n (%) | 1 (5) | 1 (5) | 0.972; 0.950 [0.055-16.293] |  |
| Previous HCV coinfection. n (%) | 4 (19) | 1 (5) | 0.199; 4.471 [0.454-44.011] |  |
| NRTI containing cART. n (%)* | 4 (19) | 14 (70) | **0.002; 0.101 [0.024-0.430]** |  |
| hsCRP (mg/L) | 0.8 [0.5-1.1] | 0.9 [0.6-1.5] | 0.324; 0.612 [0.231-1.623] |  |
| % Lymphocytes | 35.6 [25.8-38.6] | 35.2 [26.8-40.6] | 0.962; 1.002 [0.930-1.080] |  |
| % Monocytes | 5.5 [5.3-6.5] | 5.6 [5.2-6.0] | 0.518; 1.271 [0.614-2.630] |  |
| % Neutrophils | 54.3 [49.6-64.0] | 53.6 [49.2-62.5] | 0.996; 1.000 [0.938-1.066] |  |
| % CD4<sup>+</sup> Naives | 44.1 [34.5-55.6] | 44.9 [33.7-50.1] | 0.829; 0.995 [0.954-1.039] |  |
| % CD4<sup>+</sup> RTE | 78.0 [69.9-82.5] | 69.3 [64.3-76.4] | **0.016; 1.116 [1.021-1.220]** | **0.024; 1.20 [1.03-1.41]** |
| % CD4<sup>+</sup> Central Memory | 29.4 [23.3-33.2] | 28.3 [22.9-42.5] | 0.752; 0.989 [0.926-1.057] |  |
| % CD4<sup>+</sup> Effector Memory | 20.0 [18.0-27.5] | 22.0 [12.9-28.5] | 0.968; 0.999 [0.936-1.065] |  |
| % CD4<sup>+</sup> TemRA | 2.4 [1.1-4.3] | 1.6 [0.8-4.4] | 0.873; 1.017 [0.831-1.243] |  |
| % CD4<sup>+</sup> HLA-DR<sup>+</sup> | 1.8 [1.0-2.0] | 1.6 [1.1-2.3] | 0.880; 1.066 [0.464-2.451] |  |
| % CD4<sup>+</sup> ki67<sup>+</sup> | 2.4 [2.0-3.2] | 2.2 [2.1-2.6] | 0.219; 1.629 [0.748-3.547] |  |
| % CD4<sup>+</sup> CD57<sup>+</sup> | 4.8 [3.5-8.0] | 4.47 [2.1-10.8] | 0.879; 0.990 [0.873-1.124] |  |
| % CD4<sup>+</sup> CD95<sup>+</sup> | 55.0 [40.7-66.0] | 54.4 [44.1-63.9] | 0.590; 1.011 [0.971-1.052] |  |
| % CD4<sup>+</sup> CD25<sup>hi</sup> FoxP3<sup>+</sup> | 1.2 [0.9-2.0] | 1.5 [1.1-1.7] | 0.992; 0.995 [0.358-2.765] |  |
| % CD4<sup>+</sup> CD25<sup>hi</sup> FoxP3<sup>+</sup> HLA-DR<sup>+</sup> | 13.9 [9.9-21.6] | 16.1 [12.2-22.1] | 0.543; 0.973 [0.889-1.064] |  |
| % CD4<sup>+</sup> CD25<sup>hi</sup> FoxP3<sup>+</sup> ki67<sup>+</sup> | 18.5 [13.5-25.2] | 20.1 [17.6-25.0] | 0.216; 0.930 [0.830-1.043] |  |
| % CD4<sup>+</sup> CD25<sup>hi</sup> FoxP3<sup>+</sup> CD39<sup>+</sup> | 82.0 [41.7-88.2] | 83.7 [81.2-85.1] | 0.716; 1.005 [0.980-1.029] |  |
| % CD4<sup>+</sup> CD25<sup>hi</sup> FoxP3<sup>+</sup> CTLA4<sup>+</sup> | 59.0 [46.2-65.2] | 57.7 [47.8-69.2] | 0.545; 0.986 [0.941-1.032] |  |
| % mDCs | 0.5 [0.3-0.8] | 0.8 [0.6-1.4] | **0.042; 0.226 [0.054-0.949]** | **0.048; 0.16 [0.03-0.98]** |
| % pDCs | 0.2 [0.1-0.2] | 0.2 [0.1-0.3] | 0.529; 0.109 [0.000-108.297] |  |

Continuous variables are expressed as median values [IQR] and categorical variables are expressed as number of cases (&). All demographic, clinical andimmunological variables with a *p* value of <0.1 in the unadjusted model were included in the adjusted model and are shown in bold. Hence, age, CD4^+^/CD8^+^ ratio, time on HIV-treatment, &CD4^+^RTE and &mDCs were included in the enter multivariate model (n=38). Variables with a p value of <0.1 were showed in *italics*. Variables with a *p* value of <0.05 in the adjusted model were considered statistically significant and are shown in bold. *The absence of NRTIs was colineal with the presence of MVC, then, only the second variable was included in the multivariate model.

### Relationship between the time of exposure to a MVC-containing cART and immunological variables

We observed a high variability on the time of exposure to the MVC-cART previous to vaccination (median [IQR], 16 [5–38] months). Thus, we explored if this could have affected its impact on immunological variables. This analysis was logically restricted to the MVC-cART group (N=21). We observed negative correlations between the time of exposure to MVC-cART and the %CD4^+^ki67^+^ (r=-0.477, *p*=0.029), the %CD4^+^HLA-DR^+^ (r=-0.489, *p*=0.034) and the %CD4^+^CD25^hi^FoxP3^+^ (r=-0.523, *p*=0.015) (Table 3 and Figure 1).

**Table 3.** Relation-ship between the time of exposure to an MVC-containing cART and immunological variables.

| Immunonological variables (n=21) | **r** | **p** |
| --- | --- | --- |
| hsCRP (mg/L) | 0.003 | 0.989 |
| % Lymphocytes | 0.068 | 0.769 |
| % Monocytes | -0.297 | 0.191 |
| % Neutrophils | -0.079 | 0.735 |
| % CD4<sup>+</sup> Naives | 0.185 | 0.434 |
| % CD4<sup>+</sup> RTE | -0.161 | 0.497 |
| % CD4<sup>+</sup> Central Memory | -0.151 | 0.526 |
| % CD4<sup>+</sup> Effector Memory | -0.085 | 0.723 |
| % CD4<sup>+</sup> TemRA | -0.087 | 0.714 |
| % CD4<sup>+</sup> HLA-DR<sup>+</sup> | -0.489 | **0.034** |
| % CD4<sup>+</sup> ki67<sup>+</sup> | -0.477 | **0.029** |
| % CD4<sup>+</sup> CD57<sup>+</sup> | 0.110 | 0.663 |
| % CD4<sup>+</sup> CD95<sup>+</sup> | -0.170 | 0.460 |
| % CD4<sup>+</sup> CD25<sup>hi</sup>FoxP3<sup>+</sup> | -0.523 | **0.015** |
| % CD4<sup>+</sup> CD25<sup>hi</sup>FoxP3<sup>+</sup>HLA-DR<sup>+</sup> | -0.126 | 0.586 |
| % CD4<sup>+</sup> CD25<sup>hi</sup>FoxP3<sup>+</sup>ki67<sup>+</sup> | -0.318 | 0.172 |
| % CD4<sup>+</sup> CD25<sup>hi</sup>FoxP3<sup>+</sup>CD39<sup>+</sup> | 0.116 | 0.617 |
| % CD4<sup>+</sup> CD25<sup>hi</sup>FoxP3<sup>+</sup>CTLA4<sup>+</sup> | 0.226 | 0.324 |
| % mDCs | -0.133 | 0.567 |
| % pDCs | 0.105 | 0.650 |

Variables with a *p* value of <0.05 were considered statistically significant and are shown in bold.

**Figure 1.**
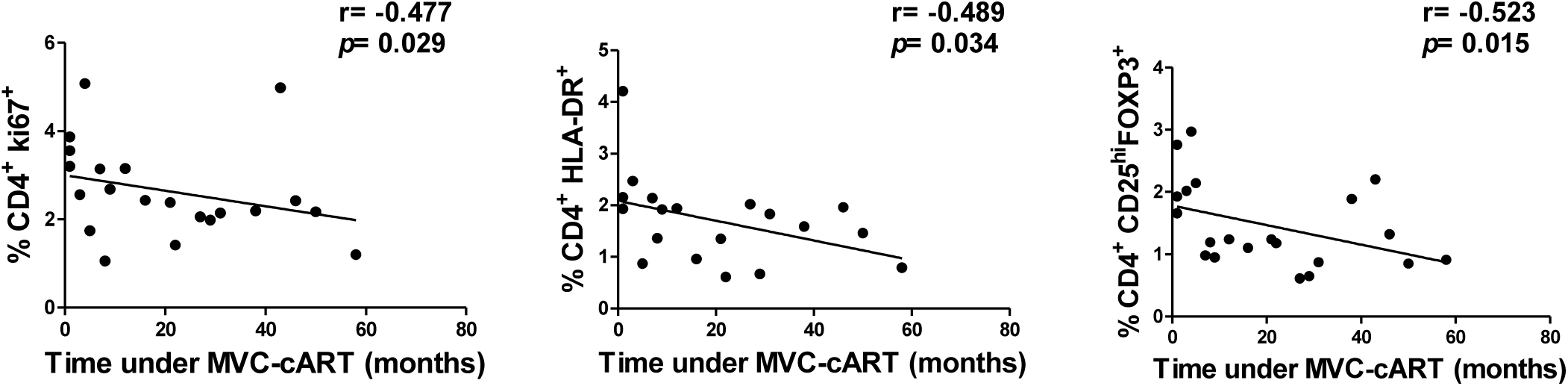
Relationship between the time of exposure to an MVC-cART and immunological variables. Only correlations with a *p* value of <0.1 between the time of exposure to an MVC-cART and different immunological variables are represented.

### Associations between the hsCRP and T-cell immunological variables

We finally explored potential correlations between the inflammation-related marker hsCRP and the main immunological variables stressed in previous analyses, in order to gain insight into both, the significance of inflammatory levels in the HBV vaccine responsiveness and the potential impact of a MVC-cART in such levels. The hsCRP inversely correlated with %CD4^+^RTE (r=-0.326; *p*=0.049), whereas positively correlated with activation-related markers as %CD4^+^CD25^hi^FoxP3^+^HLA-DR^+^ (r=0.468; *p*=0.003), %CD4^+^HLA-DR^+^ (r=0.316; *p*=0.057) and %CD4^+^CD25^hi^FoxP3^+^ki67^+^ (r=0.315; *p*=0.054), though these two last associations showing borderline significance (Figure 2).

**Figure 2.**
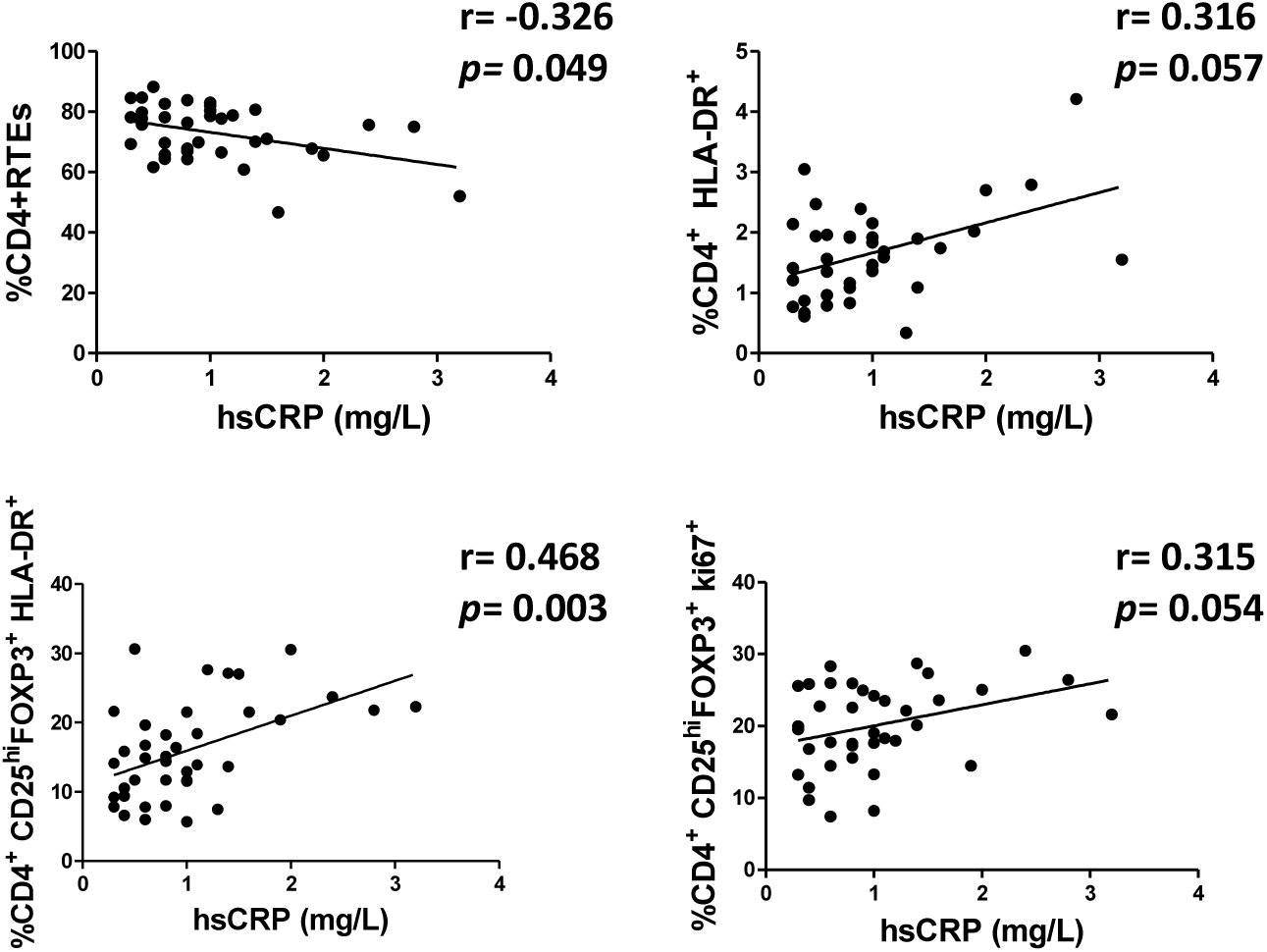
Associations between hsCRP and different T-cell immunological variables. Only correlations with a *p* value of <0.1 between hsCRP and several T-cell immunological variables are represented.

## DISCUSSION

We recently reported a beneficial effect of a MVC-cART in the response to the HBV vaccine in HIV-infected subjects younger than 50 years old [5]. We extended here this analysis and, after adjustment by baseline immunological variables, the effect of the MVC-cART remained independently associated. Additionally, a negative effect of Treg cells and a beneficial effect of pDC were also observed. Furthermore, being on a MVC-cART prior to vaccination was associated with an improved immunological profile. Such profile included lower inflammatory levels, activated CD4 T-cells and frequency of Treg and higher frequency of RTE.

In the response to the HBV vaccine, a peptide antigen administered intramuscularly, the helper CD4 T-cell function plays a major role [13, 14] and it is well assumed that T-cell exhaustion and senescence related to HIV infection may result in the failure of such response [15]. The involvement of Treg cells in several immunization models is also being studied [16, 17, 18]. The rationale is that Treg cells suppress the proliferation and cytokine secretion of CD4, CD8 T cells and monocytes, dendritic and B cells [19, 20]. In fact, Treg cells were found within germinal centers of human lymphoid tissues suppressing the B cell immunoglobulin class switching needed to mount a properly antibody response [21]. Importantly, HIV-infected subjects show higher frequency of Treg-cells in peripheral blood compared to healthy donors [22], and we already found a negative role of Treg cells in the response to HBV vaccine in a previous cohort of HIV-infected subjects [12]. Now, in this cohort, we also found the frequency of Treg, particularly those activated/proliferating (CD4^+^CD25^hi^FoxP3^+^ki67^+^), inversely associated to the magnitude of the vaccine response. Reasonably, the fraction of activated/proliferating Treg cells could be the more suppressive. We have also observed a positive association of vaccine responsiveness with the frequency of pDC. Globally, DC are a connection between the innate and adaptive immune system, playing an essential role against pathogens and during vaccination, that are being already targeted for improvement of HBV vaccine responsiveness [23, 24, 25]. Particularly, pDC are a unique DC subset that specializes in the production of type I interferons promoting antiviral responses and are also able to induce T helper 1 CD4 responses [26], what reasonably explains their contribution in the improvement on HBV vaccine responsiveness.

Also importantly, plasma levels of soluble inflammatory markers before vaccination negatively predicted responses to HAV, HBV, and tetanus vaccines in HCV and HIV infection [27]. Moreover, CRP levels were a significant predictor of herpes zoster vaccine response in elderly nursing home residents [28]. In our cohort, we observed only a borderline negative association between the hsCRP levels and the vaccine responsiveness, but these levels correlated with the frequency of CD4^+^CD25^hi^FoxP3^+^ki67^+^, a key factor in relation to the vaccine responsiveness.

Antiretroviral treatment partially restores the immunodeficiency related to HIV infection, improving antigen-specific T-cell responses and recovering T-cell repertoire [29]. In fact, the duration of cART was associated with the HBV vaccine response [30]. However, the specific effects of different antiretroviral families have been less studied. It is reasonable to expect a negative impact of nucleoside reverse-transcriptase inhibitors (NRTI), since they affect cellular senescence through inducing accelerated shortening of telomeres in peripheral T-cells [31]. In fact, telomere length has been associated to the response to influenza vaccine in elderly non HIV-infected subjects [32]. Moreover, we have recently found a better profile of T-cells, regarding biomarkers of cell survival (CD127) and replicative senescence (CD57), in subjects on NRTI-sparing regimens [33]. In contrast, MVC positively impacted the response to different vaccine antigens in HIV-infected subjects [4, 5], although the mechanisms involved in such effect have not been yet explored.

We have now explored the immunological profile associated with a MVC-cART in the context of the HBV vaccine response, including several markers of CD4 T-cell activation, senescence and susceptibility to apoptosis, as well as Treg cells and DC subsets. HIV-infected subjects with such therapy at the moment of vaccination showed better characteristics of their CD4 T-cell pool, with a less activated phenotype, a higher contribution of RTE and a lower frequency of Treg cells. Noteworthy, vaccine responsiveness was mostly associated to these factors in others cohorts. Thus, a better response has been predicted in relation with a lower activation of T-cells [34], a higher frequency of CD34+ precursors [35, 36] and a lower frequency of Treg [12]. Interestingly, we previously showed an effect of MVC in reducing Treg in antiretroviral-naïve subjects [10]. In the present cohort of cART-experienced subjects, the frequency of Treg was decreased with increasing time of exposure to a MVC-cART. Since both cohorts differed very much, not only in therapeutic parameters but also in age or time from diagnosis among other factors, this finding empowers the immunomodulatory properties of MVC through reducing Treg cells. Regarding the possible effect of MVC-cART on inflammatory levels, we failed to observe a direct association with hsCRP levels in our cohort, but we cannot exclude a potential effect of MCV-cART on other inflammatory cytokines, as previously reported [6]. Moreover, hsCRP was inversely associated with the frequency of RTE and positively associated with T-cell activation markers, suggesting also a potential indirect impact of MVC on the inflammatory state.

Ideally, we should have included more subjects to improve the power of our analyses, however, we had to restrict our analyses to the population younger than 50 years old because of the higher effect of MVC-cART on the vaccine responsiveness in this population [5]. In fact, we observed a higher impact of MVC-cART on the studied immunological parameters in this age-restricted population (data not shown). It is reasonable to speculate that the added age-associated immunodeficiency could limit or mask the potential benefits of such antiretroviral therapy on the immunological profile. Accordingly, the immunosenescence, which compromises HBV vaccine responsiveness [14], is accentuated or presents unique features in an HIV-infection scenario [37, 38]. On the other hand, it is well-known that age limits HBV vaccine responsiveness [39]. Interestingly, the group on MVC-cART showed a lower CD4/CD8 ratio, which has been reported to negatively impact vaccine response [40]. This could be due to the lower period of treatment in this group, critical for the CD4/CD8 T-cell ratio normalization [41]. In any case, despite the lower CD4/CD8 ratio, the group on MVC-cART showed better vaccine responsiveness and CD4 T-cell profile. Finally, we cannot discriminate among the particular effects due to the presence of MVC or to the absence of NRTIs in the cART and thus, we can only conclude about the beneficial effects of such combined therapy. In accordance, similar combined therapies (NRTI-free) are being explored in the clinical setting in an attempt to reduce toxicities and to improve immune reconstitution [42].

In summary, in this extended analysis, MVC-cART remained independently associated to a better HBV vaccine responsiveness in subjects younger than 50 years, but other immunological factors as Treg and pDC also showed a key role. This could focus further research on the mechanisms involved to find out novel therapeutic targets that could improve such responsiveness. We also report that MVC-cART was associated with a less activated CD4 T-cell profile, with higher levels of RTE and a lower frequency of Treg. Further research would give complementary information about the potential benefit of the instauration of similar combined antiretroviral therapies before initiating vaccination protocols with the aim of improved responsiveness.

## MATERIALS AND METHODS

### Study design, patients and samples

The vaccination protocol has been reported elsewhere [5]. Briefly, HIV-infected subjects from the Virgen del Rocío University Hospital were consecutively vaccinated against HBV. These subjects: a) were on suppressive cART (at least in the last 6 months), b) had CD4 T-cell populations of >300 cell counts/µl, c) negative serology for HBsAg and anti-HBc and d) anti-HBs titers of ≤10 mIU/ml. The vaccination protocol consisted of 3 intramuscular double doses (40 µg) of the recombinant Engerix-B vaccine (GlaxoSmithKline, Brentford, United Kingdom) at 0, 1, and 3 months. The vaccine response was measured 6 months after the first dose. A group of subjects was simultaneously vaccinated at 0 and 6 months against Hepatitis A Virus (HAV) (simultaneous HAV vaccination) with two intramuscular doses of the vaccine Havrix-1440 (GlaxoSmithKline, Brentford, United Kingdom). This subgroup of subjects had a previous negative serology for HAV. Fresh blood samples were collected at baseline, just before the administration of the first vaccine dose. All patients gave informed consent to enter the study which was approved by the Ethic Committee of our Hospital. We restricted the present analyses to subjects younger than 50 years old (n=41) from the total population because the beneficial effect of a MVC-cART on the vaccine response was observed in this population [5].

### Laboratory measurements

Absolute numbers of CD4 and CD8 T cells and percentages of lymphocytes, monocytes and neutrophils were determined with an Epics XL-MCL flow cytometer (Beckman-Coulter). Plasma HIV-1 RNA levels were measured using quantitative PCR (Cobas Ampliprep/Cobas TaqMan HIV-1 test; Roche Molecular Systems, Basel, Switzerland) with a detection limit of 20 HIV-RNA copies/ml. Plasma samples were tested for HBV-related markers (HBsAg, anti-HBs, and anti-HBc) using an HBV enzyme-linked immunosorbent assay (ELISA; Siemens Healthcare Diagnosis, Malvern, PA). Qualitative PCR amplification was used for plasma HCV amplification (Cobas Amplicor; Roche Diagnosis, Mannheim, Germany), with a detection limit of 15 IU/ml. The high sensitive C reactive protein (hsCRP) levels were determined with an immunoturbidimetric serum assay, using Cobas 701 (Roche Diagnostics, Mannheim, Germany).

### Flow cytometry

Peripheral blood mononuclear cells (PBMCs) were isolated from fresh blood before the first dose of vaccine and cryopreserved. For the immunophenotyping of cellular subsets, PBMCs were thawed and immediately stained with surface antibodies: anti-CD31 PE-CF594, anti-CD56 BV510, anti-CD25 BV605, anti-CD45RA BV650, anti-CD4 BV786, anti-CD3 APC-H7, Lin2 FITC (anti-CD3, anti-CD19, anti-CD20, anti-CD14 and anti-CD56), anti-CD11c BV650, anti-HLA-DR BV711 (BD biosciences, USA), anti-CD39 FITC, anti-CD57 PE-Cy7, anti-OX40 BV421, anti-HLA-DR BV570, anti-CD95 BV711, anti-CD27 AF700 (BioLegend, USA) and anti-CD123 AF700 (R&D, San Diego CA, USA). When necessary for intracellular staining, cells were fixed and permeabilized according to the manufacturer’s instructions (FoxP3/Transcription Factor Staining Buffer, Ebioscience, USA), and stained with intracellular antibodies: anti-ki67 PerCP-Cy5.5, anti-FoxP3 PE and anti-CTLA-4 APC (BD Biosciences, USA). Isotype controls for CD39, CD31, OX40, CD25, CD95, ki67, FoxP3 y CTLA4 were included in each experiment.

We characterized peripheral CD4 T-cells according to the distribution of their maturational subsets [naïve (CD27^+^CD45RA^+^), central memory (CD27^+^CD45RA^-^), effector memory (CD27^-^CD45RA^-^) and TemRA (CD27^-^CD45RA^+^)], also including Recent Thymic Emigrants (RTE; naïves CD31^+^) and the expression of activation markers (HLA-DR, Ki67), senescence marker (CD57) and prone-to-apoptosis marker (CD95). We also identified Treg with classical markers (CD25^hi^FoxP3^+^) and their expression of the mentioned activation markers but also of functional markers (CD39, CTLA-4). We immunophenotyped myeloid dendritic cells (mDCs) as Lin2^-^HLA-DR^+^CD123^-^CD11c^+^ and plamacytoid dendritic cells (pDCs) as Lin2^-^HLA-DR^+^CD11c^-^ CD123^+^.

Viable cells were identified using LIVE/DEAD fixable Aqua Blue Dead Cell Stain (Life Technologies, USA). One million cells of each sample were stained, and a minimum of 100,000 events of total lymphocytes and 150,000 dendritic cells were acquired. Flow cytometry was performed on a LSR Fortessa (BD Biosciences, USA). Analysis was performed using FlowJo version 9.3 (Tree Star).

### Statistical analysis

Continuous variables were expressed as medians and interquartile ranges [IQR] and categorical variables as the number of cases and percentages. Linear regressions were performed to determine factors associated with the magnitude of response (absolute anti-HBs titer). Binary logistic regressions were explored to analyze the potential impact of a MVC-cART in the clinical and immunological parameters. Variables with a *p* value <0.1 in the univariate analysis were considered in multivariable models. Correlations were assessed using the Spearman’s rho correlation coefficient. A *p* value of <0.05 was considered statistically significant. Statistical analysis was performed using SPSS software (version 22; IBM SPSS, Chicago, USA), and graphs were generated using Prism (version 5, GraphPad Software, Inc.).

## ACKNOWLEDGEMENTS

This study was funded by grants from the Fondo de Investigación Sanitaria [FIS; PI14/01693; PI16/00684], co-funded by Fondos Europeos para el Desarrollo Regional (FEDER), and the Junta de Andalucía, Consejería de Economía, Innovación, Ciencia y Empleo [Proyecto de Investigación de Excelencia; CTS2593]. The Spanish AIDS Research Network of Excellence also supported this study (RD16/0025/0019). I.H-F and I.R-S were supported by an investigator sponsored research grant from ViiV Healthcare S.L. (grant number 205644). YM.P and E.R-M. were supported by the Consejería de Salud y Bienestar Social of Junta de Andalucía through the ‘‘Nicolás Monardes’’ program [C-0013-2017 and C-0032-2017, respectively]. L.T-D. was supported by Instituto de Salud Carlos III [PFIS program; FI00/00431]. The funders had no role in study design, data collection and interpretation, or the decision to submit the work for publication.

The authors express their most sincere thanks to all the subjects included in the study and to HIV Biobank of the Spanish AIDS Research Network. We thank also Magdalena Rodriguez and Marien Gutierrez Sancho for their assistance at the Day Care Hospital (Infectious Diseases Department) and to Mª Antonia Abad and Marta de Luna for their technical assistance and to Juan Manuel Praena for statistical assistance.

## TRANSPARENCY DECLARATIONS

Authors declare no conflict of interest.

